# Coarse-Grained RNA Model for the Martini 3 Force Field

**DOI:** 10.1101/2025.04.13.648640

**Authors:** Danis Yangaliev, S. Banu Ozkan

## Abstract

In this work we developed a coarse-grained model for RNA that is compatible with the Martini 3 force field. The model is parameterized following the Martini philosophy combining the top-down and bottom-up approaches. The nonbonded interactions in the model are derived from the partitioning of nucleobases between polar and nonpolar solvents, along with calculations of the potential of mean force between bases. For bonded interactions, parameters were refined based on atomistic simulations of double-stranded RNA. Additionally, an elastic network was incorporated to maintain the structural integrity of complex RNA molecules, such as transfer RNA, and other specific RNA configurations. We present the implementation of the Martini 3 RNA model and demonstrate its ability to capture the properties of individual bases, single-stranded RNA, double-stranded RNA, and RNA−protein complexes. Compared to the Martini 2 version, the current model offers several key advantages. It is fully compatible with the updated Martini 3 force field, exhibits greater numerical stability—allowing for the successful simulation of larger RNA–protein complexes, such as ribosomes, using the standard Martini timestep of 20 fs, and it demonstrates improved agreement with all-atom models and experimental data. This new RNA model enables realistic large-scale explicit-solvent molecular dynamics simulations of complex RNA-containing systems.

**Significance:** This research introduces a new coarse-grained model for explicit water MD simulations of RNA compatible with Martini 3 software. The model demonstrates improved agreement with both all-atom simulations and experimental data, enabling more accurate and computationally efficient simulations of large RNA-protein complexes. This advancement facilitates the study of RNA and RNA-protein interactions, allowing for the modeling of larger biological complexes and paving the way for more efficient simulations of RNA systems across various fields, including therapeutics, molecular biology, and synthetic biology, where understanding RNA interactions is crucial for developing biomedical applications and advancing fundamental research.

## Introduction

Ribonucleic acid (RNA) is essential for gene expression, actively regulating protein synthesis. The importance of RNA has been appreciated since the central dogma was proposed in 1958 (1, 2). Three RNA molecules, messenger RNA (mRNA), transfer RNA (tRNA), and ribosomal RNA (rRNA) are associated with the cell’s transcription of its DNA into RNA and then translated into proteins (3). Studying RNA-protein assemblies and RNA complexes, such as the ribosome, requires computational tools that can model large length and time scales efficiently. Addressing the challenges posed by the complex and diverse structural motifs of RNA demands multi-resolution models.

Molecular dynamics (MD) simulations have evolved into a powerful technique for computational studies of protein-RNA complexes. Accurate all-atom force (AA) fields have been developed for standard biomolecular chemical building blocks such as amino acids and nucleic acids (4-11). However, atomistic molecular dynamics simulations are computationally expensive, thus a major bottleneck of atomistic molecular dynamics is the limits of accessible time and length scales. While enhanced sampling methods can extend the scope of atomistic nucleotide simulations (12), they are not universally applicable to all RNA systems and often fail to provide the needed improvement in computational efficiency.

To accelerate the conformational sampling of macromolecular assemblies and computational efficiency, system dynamics can be represented at a coarse-grained (CG) level (13, 14), where individual particles - or beads - represent groups of atoms or even entire residues rather than single atoms. Despite the main drawbacks of this approach, the reduced spatial resolution and a less accurate description of intra- and intermolecular forces, it provides significantly improved conformational sampling. The primary advantage of coarse-grained molecular dynamics (CGMD) simulations is their computational efficiency, allowing for simulations over much longer timescales, reaching up to seconds.

Coarse-grained models for RNA have been developed with varying levels, depending on the specific goals of the modeling study. These models simplify the representation of RNA nucleotides by describing each nucleotide with a limited number of particles or pseudo-atoms, striking a balance between computational efficiency and the level of structural detail required. Some of the earliest and most coarse CG models represent each RNA nucleotide with just one, two or three (15, 16) pseudo-atoms or CG beads. One of the newer models even represents 3 nucleotides with a single particle and is parametrized using a well-established nearest-neighbor model (17). These highly simplified models typically rely on experimental data as external constraints to enhance the accuracy of structural predictions, focusing on capturing the general architecture of RNA molecules rather than detailed dynamics or interactions. In recent years, advancements in computational power and a deeper understanding of RNA behavior have led to the development of less coarse, higher-resolution CG models. Examples include the HiRE-RNA model by Pasquale and Derreumaux (18), the SimRNA model by Boniecki et al. (19), the oxRNA model by Šulc and co-workers (20), IsRNA model by Zhang and Chen (31) and the model introduced by Ren and co-workers (22). These models increase the granularity by incorporating a more detailed representation of the RNA backbone and/or nucleotide bases. Such enhancements significantly improve the ability to simulate RNA behavior under various conditions. These higher-resolution models enable realistic simulations of complex processes such as RNA hybridization, where two complementary strands of RNA form a duplex; RNA supercoiling, which involves the twisting and folding of RNA strands; and RNA quadruplex formation, a structure involving guanine-rich sequences. By capturing finer structural and dynamic details, these models provide deeper insights into RNA mechanisms, offering greater predictive power for RNA folding pathways, interaction networks, and functional dynamics in biological systems.

Although many specialized RNA models perform well in their targeted applications, there remains a need for a CG RNA model that can be integrated into complex biological systems. Currently, there is no single, widely accepted setup for CGMD simulations, so each study typically tailors its model to meet specific requirements—such as using implicit solvent RNA-protein models (23, 24) or models representing proteins solely by their Cα atoms (25). One promising approach capable of large-scale, explicit-solvent molecular dynamics simulations of complex RNA systems is the Martini DNA/RNA model (26, 27). It, however, is outdated and not directly compatible with its latest updated model for protein–protein and protein–lipid interactions, Martini 3 (28). Therefore, we have developed an RNA model within the Martini 3 force field framework that can be used in combination with other Martini models, such as the existing models for proteins, carbohydrates, and lipids at a wide range of solvent conditions.

A widely used Martini force field is a CG model that was originally derived for CGMD simulations of biological membranes (29). Later, it was extended to also include proteins (30) and nucleic acids (26, 27). Its implementation in widely used MD simulation software such as GROMACS makes its application very straightforward. In the Martini model, four heavy atoms plus associated hydrogens are represented usually by a single CG bead. However, small ring-like fragments or molecules (e.g., the nucleobases) are typically mapped at higher resolution. Bonded interactions between CG beads are modeled using standard interaction potentials for covalent bonds, bond angles, and dihedral rotations. Nonbonded interactions between neutral beads are exclusively described by Lennard-Jones potentials, while charged beads also include Coulombic interactions. In the parameterization process of the Martini model, top-down and bottom-up approaches are combined. The top-down parametrization of the force field is based on free energies. The main targets are experimental data like densities of liquids and transfer free energies of small solutes partitioning between polar and nonpolar solvents, which are used to determine nonbonded interaction parameters. The bottom-up parametrization relies on atomistic reference simulations, which are primarily used to extract bonded interaction parameters. A detailed description of the Martini force field can be found in (28).

Here, we developed the RNA parameters according to the general strategy for Martini parameterization, i.e., combining a top-down and bottom-up approach. The mapping and bead type selection were systematically parameterized according to Martini philosophy and, therefore, compatible with other Martini models for biomolecules and solvents. The bonded interactions were fitted to bond, angle, and dihedral distributions derived from atomistic simulations of dsRNA. The performance CG models for ssRNA and dsRNA was evaluated against all-atom models, with a focus on local and global flexibility. Additionally, we tested the stability and reliability of the new RNA parameters in various protein-RNA complexes, including ribosome.

## Results and Discussion

The parameterization approach we take in this work follows the general strategy for Martini and is similar to the one J. Uusitalo et al. used for the Martini 2 version of RNA/DNA force field (27). However, we implemented significant changes to enhance the model’s numerical stability and overall performance, while ensuring compatibility with Martini 3. Additionally, despite the introduction of smaller bead sizes (S and T) in Martini 3 (28), we had to incorporate new bead types into our model to achieve improved base packing. In a dsRNA, bases are stacked very closely, with an inter-base distance of only 0.34 nm—still too small for tiny (T) beads alone. The new bead types are derived from standard Martini 3 beads and maintain full compatibility when interacting with them, but they have reduced van der Waals radii for interactions with each other.

### Corse-Grained Mapping

The first step in the Martini 3 parametrization procedure involves virtually fragmenting the molecules and assigning them to appropriate bead types. The mapping process determines which atoms at the all-atom level are represented by a single bead in the CG model. Typically, within the context of Martini 3, one can map two to six heavy atoms into a single bead. The number of atoms mapped to each bead, together with the degree of branching, dictates the bead sizes, and it is generally advisable to keep functional groups intact as much as possible to preserve their chemical identity and interactions. Each bead size in the Martini model has four main types of particles: polar (P), intermediate-polar (N), nonpolar (C), and charged (Q), with 6 subtypes dictating the degree of polarity of a bead. All these parameters together will determine the strength of non-bonded interactions between the particles.

Figure 1 illustrates the chosen mapping and bead assignments for the nucleotides involved in this study. For each residue, the beads are divided into backbone beads (BB1, BB2, and BB3) and side chain beads (SC1 through SC6). It’s worth noting that there are several major differences compared to the previous model of Martini 2 (26, 27). The most significant change is the revised backbone mapping, motivated by two main factors. First, the three-bead backbone of the previous version is a major source of numerical instability due to the bead alignment when calculating dihedral angles involving these beads. As described by Bulacu et al. (31), traditional torsion angle potentials (i.e., proper dihedrals) used in atomistic MD simulations may not always be suitable for modeling CG systems (32). Second, the new mapping better preserves the inherent symmetry of the RNA backbone, where the phosphate group and the C3’ carbon form a saw-like pattern. In the new model, the first backbone bead (BB1) still models the phosphate group, but the 3′ end of the sugar moved to the second bead (BB2). The hydroxyl group in the ribose moves the third bead (BB3).

**Figure 1.**
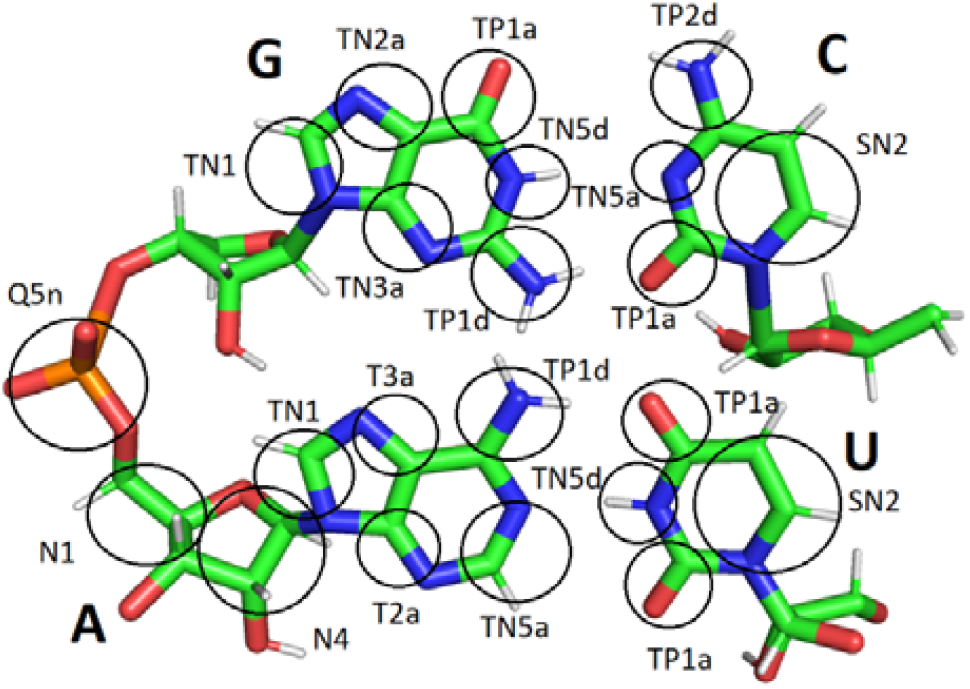
Coarse-grained mapping for the model. The RNA backbone is modeled with one bead describing the phosphate and two beads describing the sugar. The pyrimidines are modeled with four beads, adenine with five and guanine with six. The Martini bead type of each bead is shown.

One of the major changes in the Martini 3 model was the introduction of small (S) and tiny (T) beads, which can effectively model the smaller, more intricate segments of the molecules and are better suited to represent ring-like compounds. In this work we utilized them to model the small and rigid rings of pyrimidines and purines. Cytosine and uracil are now mapped onto four tiny beads, adenine onto five, and guanine—six. We also took advantage of the newly introduced hydrogen bonding labels. Even though Martini does not have any special interactions to model directional hydrogen bonds, which are crucial for the formation of dsRNA and the specificity of base pairing, hydrogen donor (“d”) and acceptor (“a”) character labels can be added to all P and N beads to better describe the specificity of base pairing.

### Parametrization of Bonded Terms

Bonded interactions are typically obtained via a “bottom-up” approach based on reference atomistic MD simulations. The aim is to match the CG conformations available in Martini as closely as possible to the conformational space of the reference AA model; we used CHARMM36 in our work (33, 34). Martini employs simple bonded potentials, including two-body harmonic, three-body angular, and four-body periodic dihedral potentials, whose parameters are derived from atomistic simulations mapped onto a CG trajectory. In this work, however, we slightly altered this approach. In contrast to the previous version, where most of the non-bonded interactions within a residue were turned off, we decided to exclude only the non-bonded interactions between the nucleoside beads (SC1-SC6) and follow the standard Martini exclusion rule where non-bonded interactions between connected beads are always excluded. This, however, led to a problem: when we tried to use the bonded parameters obtained from matching them with distributions from short-length single-stranded RNAs, in some cases it led to highly unstable conformations of bigger molecules like tRNA due to steric clashes within the structure because of the very tight packing of double-stranded RNA and the mismatch between the VDW radii of atoms and martini beads. To alleviate that, instead of using only ssRNA for the parametrization procedure, we used the double-stranded, and every few steps of the iterative matching procedure we added a step during which we removed the constraints between the side chain beads, froze the backbone, added position restraints to the sidechains, and ran energy minimization, which helped to remove the steric clashes while slightly altering the bond lengths of the side chains. As a result, some of the obtained coarse-grained distributions are slightly shifted with respect to their atomic counterparts. To improve the agreement between CG and AA equilibrium distances, we replaced the non-bonded interactions between specific bead pairs, namely BB3 and BB1 of the subsequent residue, and BB2 and SC2, with a Morse bond-stretching potential. This modification allowed for a more accurate representation of equilibrium distances as observed in the reference AA distributions. The complete list of bonded terms incorporated into the CG model is detailed in Figure S1 of the Supporting Information. Examples of the corresponding AA and CG for the backbone bond distributions are shown in Figure 2.

**Figure 2.**
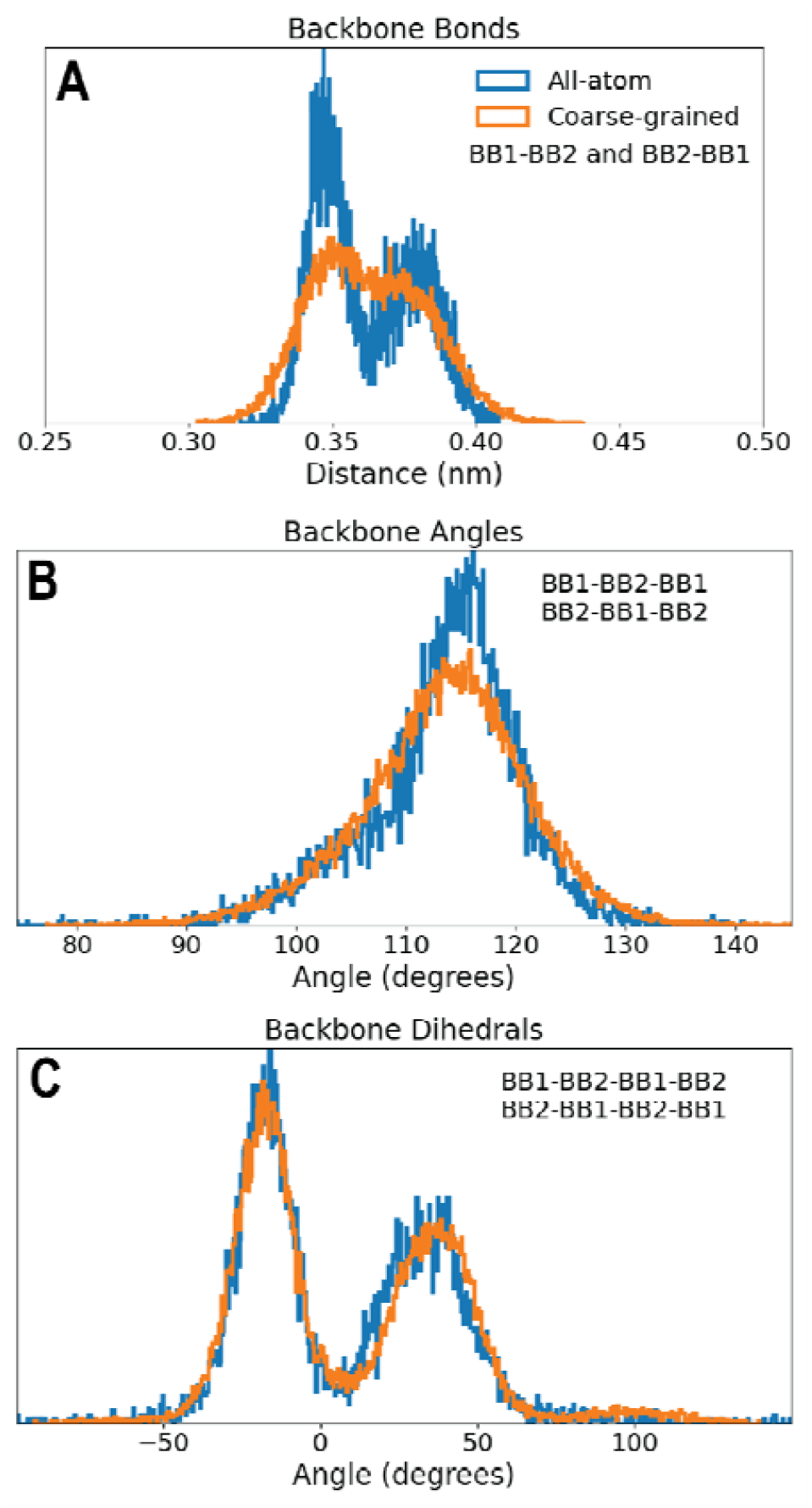
Example distributions of bonded terms. Bond (A), angle (B), and dihedral angle (C) distributions for consecutive backbone beads in the CG Martini model (orange) and AA CHARMM force field (blue). The CG distributions are smoother and broader than the AA distributions, reflecting the reduced resolution of the CG model. This softness in CG bonds is an intentional tradeoff to enhance the model’s applicability for simulating complex tertiary RNA structures, such as tRNAs, while maintaining numerical stability. A comprehensive comparison of AA and CG distributions across all bonded terms is provided in Figure S1 of the Supporting Information.

While both AA and CG distributions display similar overall trends, there are notable distinctions. As expected for a CG model, the CG distributions are generally smoother, reflecting the reduced resolution inherent to coarse graining. Additionally, the bimodality observed in some AA distributions is not captured by the CG model, as such fine-grained details are beyond the scope of coarse-grained representations. We also had to make tradeoffs to improve the numerical stability of the model compared to the Martini 2 version. One such adjustment is the rigidity of bonds within nucleobases, which we modeled using a combination of constraints and virtual sites. While this approach results in excessively stiff bonds, it significantly improves numerical stability. On the other hand, some backbone potentials in the current model are intentionally softer than their all-atom counterparts, allowing the simulation of more complex molecules, beyond just single- or double-stranded RNA. Overall, these modifications strike a balance between maintaining computational efficiency, ensuring numerical stability, and preserving key structural and dynamic features of RNA molecules, thus broadening the utility of the CG model for a wider range of applications in RNA biophysics.

### Partitioning Free Energies

The Martini force field is designed to reproduce free energies of transfer between different solvents, making the partitioning of building blocks between polar and nonpolar environments a critical benchmark test for bead type assignment. In this study, the selection of bead types for the nucleotide bases were selected to match the experimental free energies of transfer from water to octanol for the bases and from water to chloroform and water to 2-butanol for their methylated analogs. Coarse-grained (CG) free energy values were systematically compared to available experimental data (35, 36) to validate and refine the model.

The initial bead assignments were done following the standard Martini protocol and were subsequently adjusted based on partial charge distributions derived from atomistic OPLS-AA force field parameters (37). This refinement was necessary because groups of atoms with identical structures may exhibit varying degrees of polarity and different behaviors depending on their molecular context. For example, in adenine, beads SC2 and SC5, despite being structurally identical, were assigned different types to reflect their distinct polarities. We then proceed to iteratively adjust the bead types to achieve an optimal agreement with reference data across all three solvents. The final partitioning free energies, presented in Figure 3, demonstrate the effectiveness of the refined RNA model. The current model achieves good agreement with experimental values, with a mean absolute error of less than 2 kJ/mol, well within the acceptable range for Martini beads (28). This represents a significant improvement over the previous model, thanks to the increased variety of bead types in Martini 3, highlighting the importance of carefully tuning bead types to capture the unique chemical environments of RNA nucleotides.

**Figure 3.**
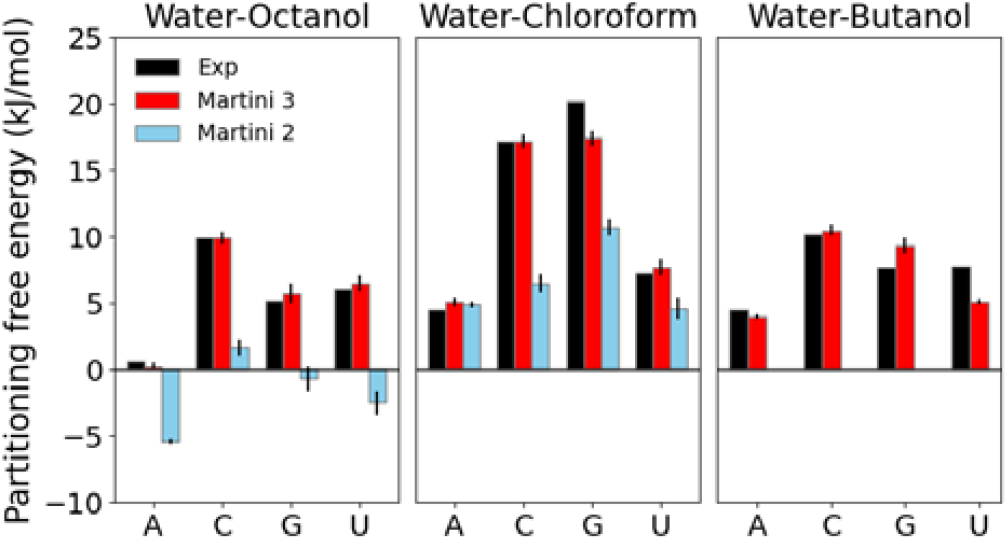
Partitioning free energies of RNA nucleobases (A, C, G, U) between water and non-polar solvents: water-octanol, water-chloroform, and water-butanol. Experimental data (black) are taken from Ref. 49 and 50, Martini 3 data (red) represent the current model, and Martini 2 data (cyan) are reproduced from Ref. 23. The Martini 3 model demonstrates improved agreement with experimental measurements compared to Martini 2, particularly in capturing the relative partitioning trends across different solvents.

### Dimerization and Hydrogen Bonding of Nucleobases

The partitioning simulations provided insights on how the nucleobases interact with their environment. Next, we examined how these bases interact with each other in water. The key aspect of DNA and RNA annealing is the specificity of base pairing, which is largely governed by hydrogen bonding. These interactions play a crucial role in ensuring complementarity during strand annealing. To investigate this, we constrained the bases to remain within a plane and calculated the Potential of Mean Force (PMF) as they were brought closer together along a defined axis. Figure 4 illustrates representative PMF profiles, while the full set of profiles for all nucleobase pair combinations is provided in Figure S2 of the Supporting Information.

**Figure 4.**
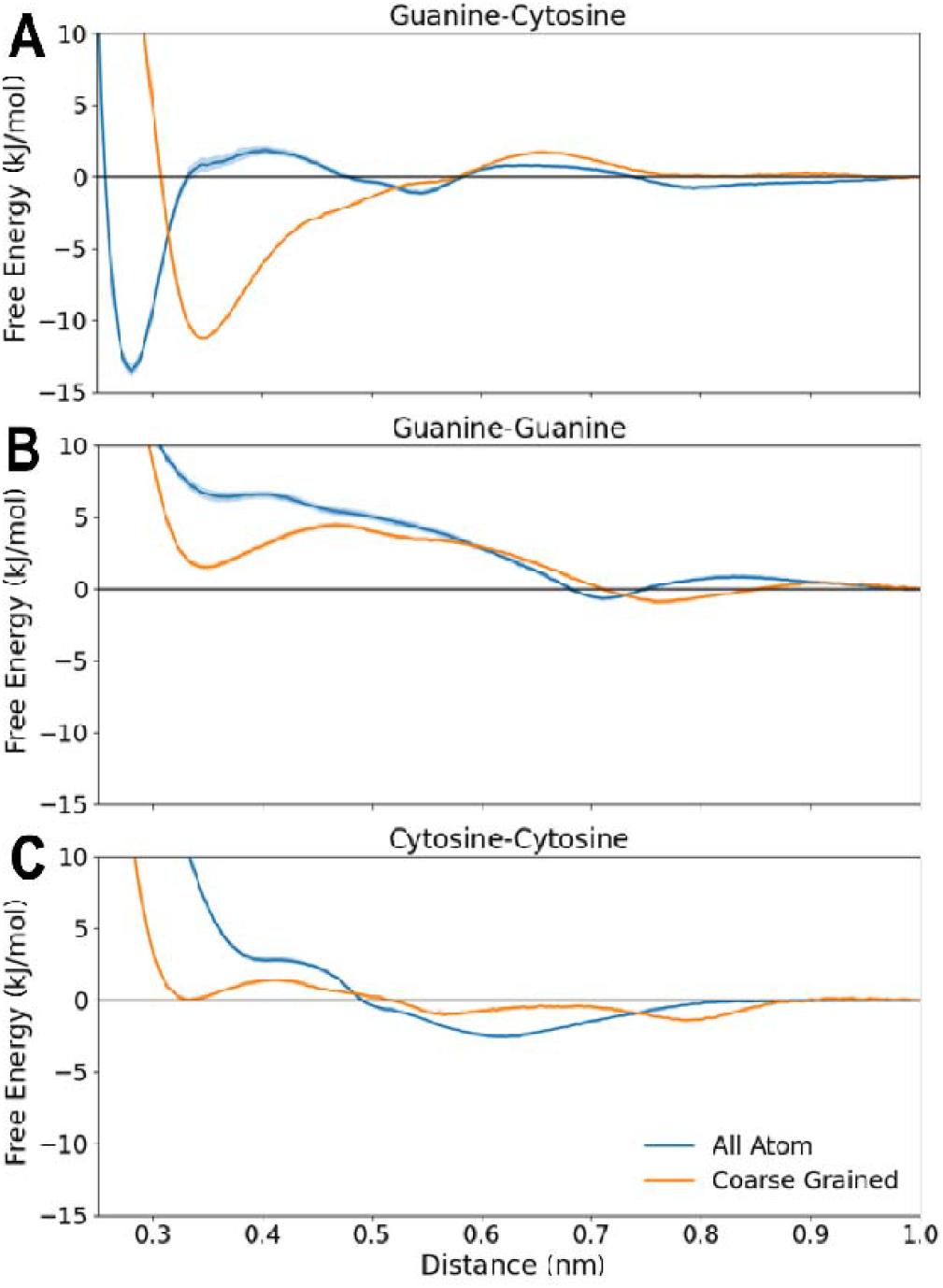
Base-base interactions. PMF profiles for interactions between guanine and cytosine (A), guanine and guanine (B), and cytosine and cytosine (C) under planar constraints. In these simulations, the bases were restricted to remain in a plane and subsequently subjected to pulling along their separation distance to characterize hydrogen-bonding interactions. PMF curves are presented for the OPLS all-atom force field (blue) and the Martini coarse-grained force field (orange). For clarity, error bars corresponding to the Martini force field are too small to be discernible. The results for other nucleotide base pairs can be found in Figure S2 of the Supporting Information. These comparisons highlight the differences in interaction strength and distance-dependent profiles between atomistic and coarse-grained models.

To better represent hydrogen bonding interactions in the coarse-grained model without significantly altering stacking interactions, we introduced new bead types. This adjustment was necessary because the spherically symmetric nature of coarse-grained potentials links hydrogen bonding and stacking interactions, making it challenging to modify one without affecting the other. Specifically, we slightly increased the interaction strength for beads forming hydrogen bonds to better replicate the all-atom PMF profiles while reducing it for beads that do not engage in hydrogen bonding.

Despite these refinements, discrepancies remain between the all-atom and coarse-grained PMF profiles. While the coarse-grained and all-atom profiles match qualitatively, the free energy minima in the coarse-grained PMF profiles are shifted relative to those in the all-atom profiles (Figure 4). This discrepancy arises because the size of the coarse-grained beads is based on the distance between two stacking bases, and the increased hydrogen bonding distance cannot be fully compensated by adjusting the bead size. Furthermore, coarse-grained beads are mapped to the geometric centers of the underlying atoms, resulting in a larger effective separation between nucleobases compared to the atomistic hydrogen-bonding distances.

To address the above limitations, we plan to introduce a Go-like network (38) or improve electrostatics parametrization in future iterations of the force field. This approach will provide a more accurate description of hydrogen bonding interactions by targeting base-base stacking interactions as the primary parameterization objective. By incorporating this refinement, we aim to improve the overall accuracy of the coarse-grained model for both hydrogen bonding and stacking interactions between nucleobases.

### Lipid Membrane Partitioning

In the Martini 2 model (27), the general features of the AA and CG potential of mean force profiles for moving nucleobases across a lipid bilayer matched well for all tested systems. However, for cytosine, the central barrier was lower in the CG simulations. This discrepancy might be due to water penetration into the membrane on the molecular scale (39), which is limited in the coarse-grained representation. In Martini 2, water beads are constructed to represent four water molecules, which may not fully capture the detailed dynamics of water interactions.

To address this, we assessed whether the expanded variety of bead sizes in Martini 3 could improve the agreement between the AA and CG models. Specifically, we used a smaller bead size representing two water molecules (TW) instead of the standard size (W). Figure 5 illustrates the PMF profiles for cytosine translocation toward the center of a POPC bilayer along the axis perpendicular to the membrane, comparing AA and CG simulations. The overall features of the PMF profiles in both models were similar but the use of tiny water beads in CG simulations did not result in significant improvement compared to the regular water beads.

**Figure 5.**
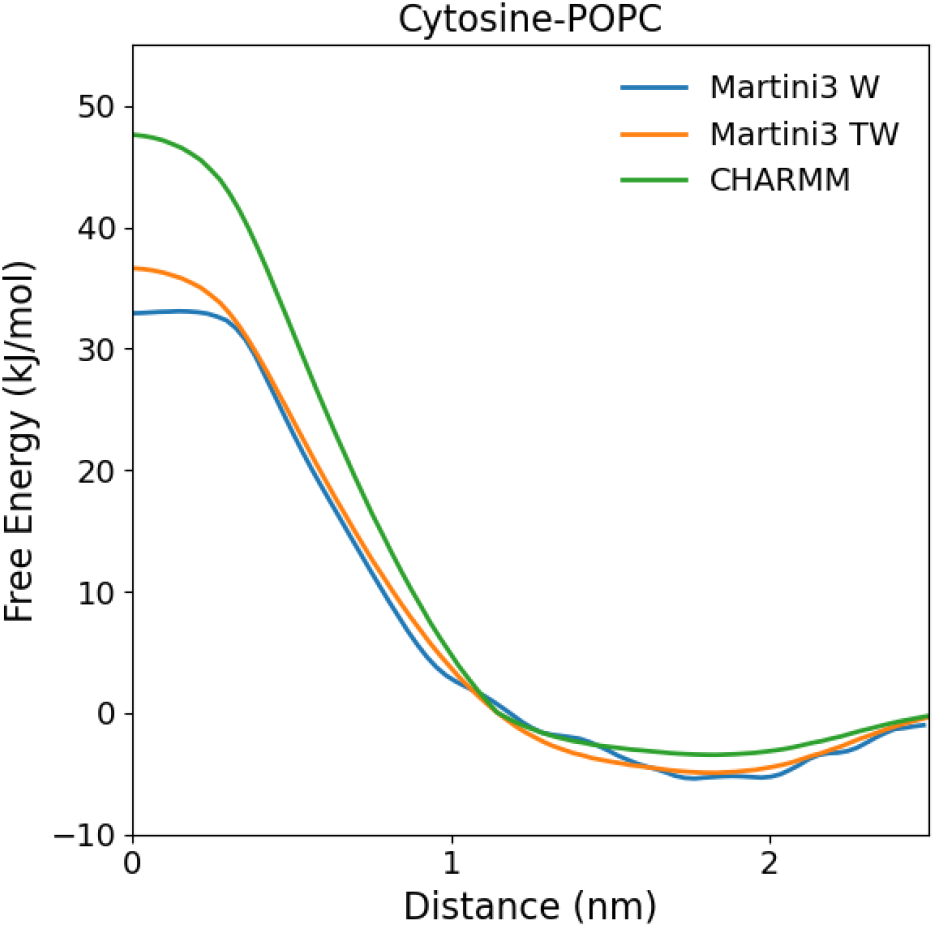
Transfer of cytosine into a lipid bilayer. PMF profiles of Cytosine transitioning away from the middle of a POPC bilayer, shown for CHARMM and Martini with both regular and tiny water bead sizes (green, blue, and orange lines, respectively).

Both AA and CG simulations show a distinct free energy barrier at the bilayer center, which is attributed to the polar nucleobases being drawn into the hydrophobic tail region of the lipid bilayer. The free energy value drops sharply as the nucleobases move away from the bilayer center, maintaining a comparable gradient in both AA and CG simulations. The minimum of free energy is observed at the interface between the hydrophobic lipid tails and the hydrophilic lipid headgroups. This reflects the amphipathic properties of the nucleobases and their preferential localization near the interface.

### Persistence Length Analysis and ssRNA folding

A commonly used measure of the large-scale properties of a polymer is its persistence length (40). A general definition is given by

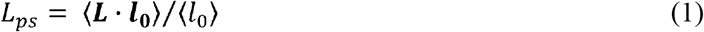

With ***L*** being the end-to-end vector of the polymer and ***l***_0_ representing the vector between the first two monomers. For an infinitely long, semiflexible polymer in which the correlations in alignment decay exponentially with separation, Eq.1 is equivalent to the commonly used form:

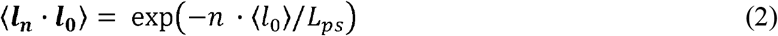

To measure the persistence length, we simulated the double-stranded RNA sequence of 100 bp length that Uusitalo et al. used in their Martini 2 work. The simulations were run in 100 mM NaCl solution without elastic network for 10 microseconds. To obtain persistence length values that are valid for long strands where end effects are negligible, the behavior of bases near the end of strands was ignored. The correlation of the helix axis (defined as the distance between consecutive base-pair midpoints) at two points was observed to decay exponentially with distance (Figure 6), allowing an estimate of *L*_*ps*_ through Eq. 2. We find a model persistence length of around 260 base pairs, somewhat stiffer but in reasonable agreement with experiment (41, 42).

**Figure 6.**
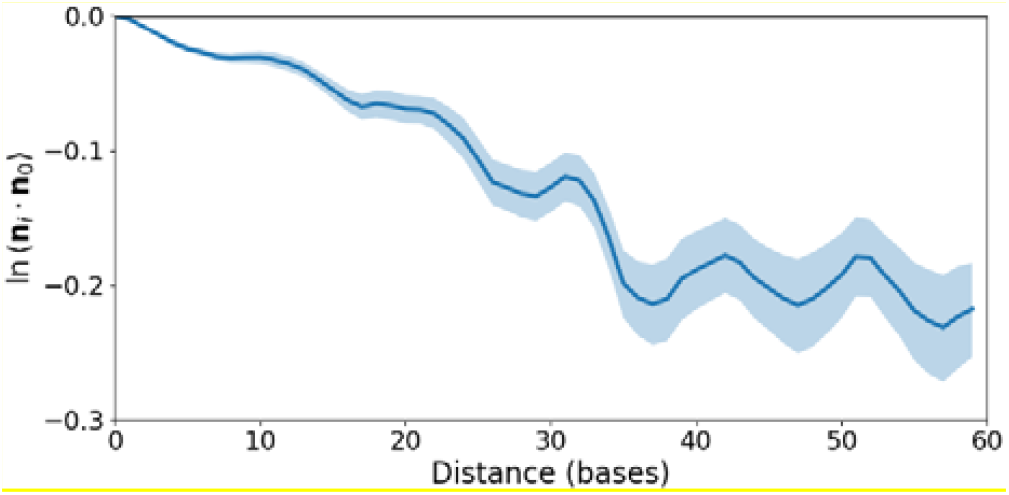
Persistence length of dsRNA. A correlation plot illustrates the determination of the persistence length. The y-axis represents the average angle ⟨***n***_*i*_ · ***n***^0^⟩ between vectors, where each vector connects the centers of two consecutive bases. The x-axis shows the distance (in bases) between these vectors.

Beyond capturing the mechanical properties of double-stranded RNA, we also tested the model’s ability to describe ssRNA folding and qualitatively reproduce all-atom folding dynamics (See Supplemental Movie 1). Despite the good performance of the model with dsRNA, its ability to describe tertiary RNA structures is still limited without using an elastic network to stabilize the structure. This is expected since we parametrized the model using dsRNA as the reference. We anticipate that incorporating a dependence of bonded parameters on secondary structure or more detailed and comprehensive fitting of backbone dihedrals will improve the model’s ability to capture RNA folding in future versions.

### Dynamics of RNA-Protein Complexes

The primary objective of this work was to develop a Martini 3-compatible force field capable of simulation equilibrium dynamics of large protein-RNA complexes. To evaluate the compatibility of the Martini RNA force field with other biomolecules, we performed simulations of a 70S ribosome and compared the results to available atomistic data (43). The elastic network was used for the RNA and Go-Martini model (44) for the proteins. The force field was shown to be numerically stable with the standard Martini 20-fs timestep with or without elastic network. Although the elastic network used in the simulations was deliberately soft (k = 100 kJ/mol/nm^2^) our model could reasonably capture the flexibility of the 5S, 16S and 23S ribosomal RNA. Figure 7 compares the typical backbone RMSF profiles of rRNA and proteins between all-atom and coarse-grained simulations. Notably, the overall RMSF values for both proteins and RNA are in good agreement with the all-atom data showing that the CG model can reproduce the flexibility of RNA backbone with complex tertiary structures and in close contact with proteins. In agreement with the results of the previous section, the CG RNA backbone is slightly stiffer than all-atom due to the lower-than-normal base-base stacking energies resulting from the chosen parametrization of hydrogen bonding and the spherical nature of Martini interactions. As we discussed above, using a different hydrogen-bonding parametrization strategy will result in better agreement with the atomistic models.

**Figure 7.**
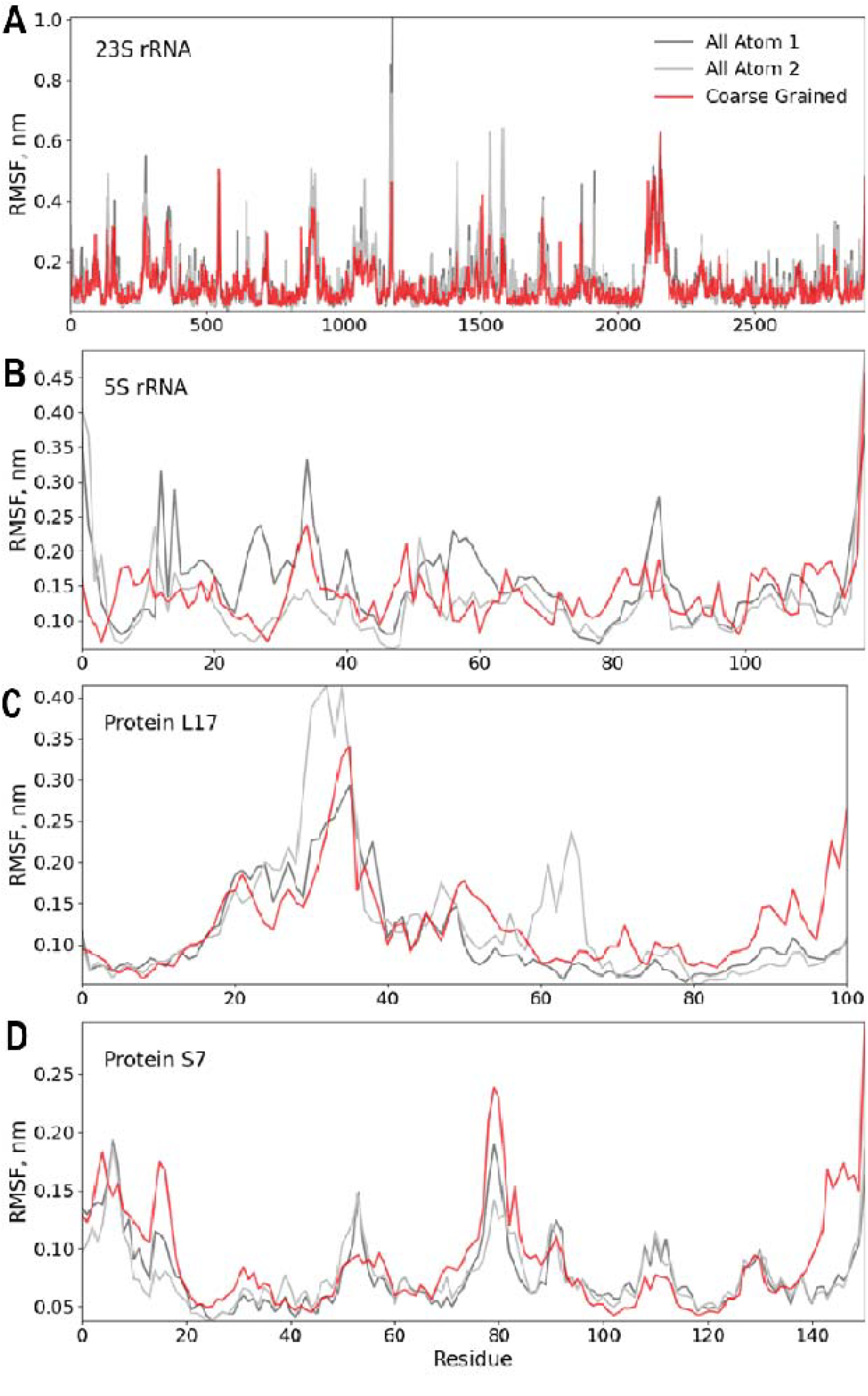
RMSF of rRNA and proteins in a 70S ribosome. Comparison between CG and AA RMSF for rRNA and two proteins that are in direct contact with it after 2 microseconds of MD simulations. CG martini is shown in red, and 2 AA simulations are in grey and silver. (A) 23S ribosomal RNA. (B) 5S ribosomal RNA. (C) Large subunit protein L17. (D) Small subunit protein S7.

To further assess the model’s ability to describe RNA-protein complexes, we investigated whether it could capture allosteric coupling in these systems. Specifically, we compared our model, along with Martini 3 for the protein, against the elastic network model (ENM), a coarse-grained approach that represents proteins as beads connected by springs. ENM has been effectively utilized in RNA-protein systems, including studies on ribosome conformational dynamics (45) and allostery-driven viral capsid assembly (46). In this study, we analyzed a eukaryotic ADAT2/3 deaminase bound to a full-length tRNA (PDB Entry: 8AW3, Figure S3), an enzyme that catalyzes the deamination of adenosine to inosine at the ‘wobble’ position of the anticodon of tRNAs (47, 48). This enzyme is a heterodimer consisting of two subunits: ADAT2, which contains the catalytic site responsible for converting adenosine to inosine, and ADAT3, which primarily plays a structural and substrate-recognition role. In the structure, the full-length tRNA adopts its characteristic L-shaped conformation, with the anticodon loop positioned into the active site of ADAT2.

To quantify coupling between the residues, we obtained backbone beads position-position cross-correlations from ENM and Martini 3 simulations. For ENM we connected all alpha carbons (Cα) of the proteins and phosphate/ribose sugar backbone atoms (P and C1’) of RNA. For Martini, separate elastic networks—with a spring constant of k = 500 □ kJ□mol^−1^□nm^−2^—were applied to each protein monomer and the tRNA. Comparison of the heatmaps, which represent the magnitude of correlation between each pair of beads (Figure 8), shows that both models exhibit similar coupling patterns for the protein and RNA domains. While a detailed analysis is beyond the scope of this study, some differences can be attributed to disordered protein regions, where ENM significantly underestimates fluctuations. Our findings demonstrate that both models predict similar allosteric coupling patterns in ADAT2/3-tRNA complex, suggesting they are effective in describing RNA-protein interactions. This consistency highlights the potential of our Martini RNA model for studying allosteric mechanisms in other RNA-protein complexes.

**Figure 8.**
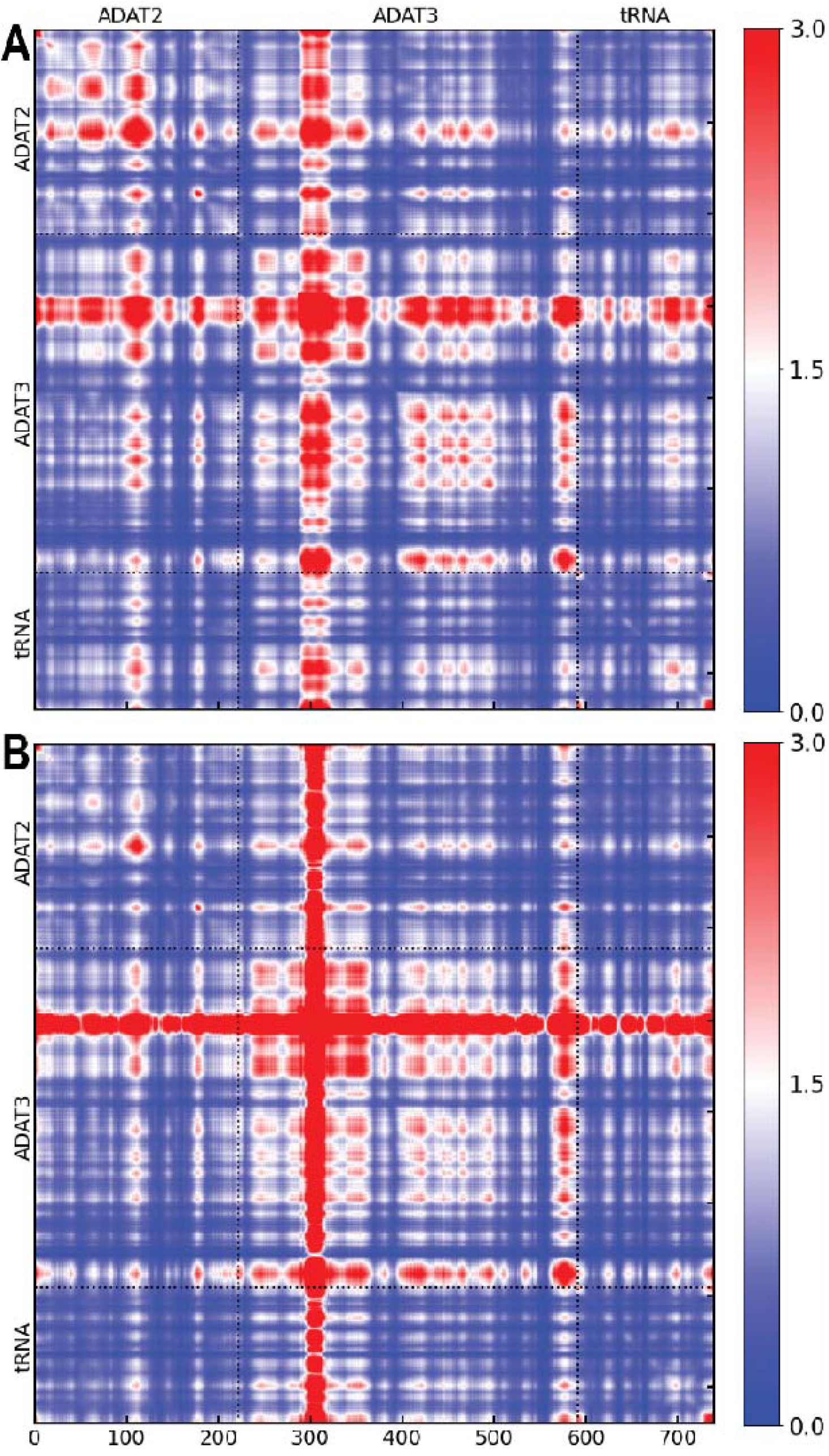
Correlation analysis of ADAT2/3 deaminase bound to a full-length tRNA. (A) Martini model. (B) ENM. The heatmap values are normalized so that the overall average is equal to 1.0 with 0.0 corresponding to uncorrelated motions. The Martini model shows some loss of detail compared to the ENM, partly due to the high flexibility of intrinsically disordered protein regions, which are not fully captured by the ENM.

## Conclusions

In this work, we have developed a coarse-grained RNA model that is fully compatible with the Martini 3 force field, leveraging the parameterization framework established in previous Martini models. This new model offers significant advancements, including improved numerical stability, enhanced agreement with experimental and atomistic data, and the ability to simulate large and complex RNA structures, such as ribosomes and tRNA-bound ADAT2/3 deaminase.

The expanded variety of beads in Martini 3 allowed for improved accuracy in RNA base stacking and hydrogen bonding by calibrating nucleobase interactions to match experimental partitioning free energies and atomistic PMF profiles. Additionally, our modifications to the RNA backbone structure have led to a more stable representation of double-stranded RNA while maintaining compatibility with other Martini biomolecular models.

To assess the model’s capability in simulating large RNA-protein complexes, we conducted CG molecular dynamics simulations of the 70S ribosome and ADAT2/3. The ribosome simulations demonstrated that the model successfully captures RNA backbone flexibility and global conformational dynamics while maintaining numerical stability, even with the standard Martini 20-fs timestep. Additionally, the model’s ability to reproduce atomistic root-mean-square fluctuations (RMSF) suggests that it can accurately represent RNA-protein interactions in large assemblies.

The ADAT2/3 simulations further validated the model’s potential in studying allosteric coupling within RNA-protein complexes. Our correlation analysis revealed that the coarse-grained model can replicate RNA-induced allosteric effects observed in other coarse-grained approaches. The agreement between our Martini-based approach and the ENM highlights the model’s ability to capture key functional dynamics in RNA-protein interactions.

Despite these improvements, certain limitations remain. The Martini model’s inability to capture directional hydrogen bonding restricts its application in studying processes such as hybridization, melting, and intercalation. Future work will focus on refining electrostatics, which may require incorporating higher-resolution elements to address this issue. Moreover, while the current model is optimized for dsRNA, further refinements are needed to accurately describe single-stranded RNA and molecules with complex secondary structures, such as tRNA.

Nonetheless, the model’s robustness in simulating large biomolecular assemblies makes it highly suitable for studying RNA–protein interactions, RNA–RNA interactions, and RNA–lipid interactions in biologically relevant environments. The successful application of this model to ribosome and ADAT2/3 simulations highlights its potential for exploring complex macromolecular dynamics, including large-scale allosteric regulation and RNA-mediated conformational changes.

Moving forward, future developments will aim to improve hydrogen bonding specificity, refine backbone flexibility for ssRNA applications, and expand the model’s applicability to a broader range of RNA structures.

## Methods

### Molecular Dynamics Simulations

MD simulations were performed using GROMACS version 2023 (49, 50). For the CGMD simulations of ssRNA, dsRNA and tRNA-bound ADAT2/3 deaminase, the molecular systems were energy minimized for 5,000 steps using the steepest descent algorithm and then equilibrated at room temperature for 100Cns with a 20-fs integration time step with position restraints on the RNA/protein backbone (k = 500□kJ□mol^−1□^nm^−2^) and 100Cns without restraints. Ribosome simulations underwent a more extensive protocol, including a 20,000-step energy minimization, a 10-ns heating phase to 300 K with a 10-fs timestep, and a 100-ns equilibration phase with a 10-fs timestep under backbone position restraints. This was followed by 1 μs of equilibration without position restraints using a 20-fs timestep. Periodic boundary conditions were applied in all cases.

During the equilibration and production stage, the temperature was maintained constant at 300□K using a modified Berendsen thermostat (51) with a τ_*T*_ coupling constant of 1□ps. The pressure was maintained at 1 bar (isotropic coupling) using a modified Berendsen barostat (52) during the initial equilibration (τ_*P*_□= □4□ps, compressibility of 3.4□× □10^−4^Cbar^−1^) and a Parrinello-Rahman barostat (53) (τ_*P*_□= □12□ps, compressibility of 3.4□× □10^−4^Cbar^−1^) for the second equilibration and all production runs. Electrostatic interactions were cut off at 1.1 nm using the reaction-field method (54), and Lennard-Jones interactions were cut off at 1.1 nm using the potential-shift Verlet method.

For atomistic reference simulations, CHARMM-GUI (55-56) was used to generate GROMACS input files, and simulations were performed using the CHARMM36 force field (33, 34) under the corresponding settings.

### Transfer Free Energy

Thermodynamic integration (58) was used to calculate the free energies of solvation. The solute was solvated in a pre-equilibrated solvent box with dimensions of 5 × 5 × 5 nm^3^. Four solvents were analyzed: pure water, pure chloroform, pure 2-butanol, and water-saturated octanol with a 0.25:0.75 water/octanol molar ratio (59). To calculate each CG free energy value, 11 simulations with uniformly distributed λ parameters were performed. In these simulations, solute-solvent interactions were progressively scaled from full interactions (λ=1) to no solute-solvent interactions (λ=0). Each simulation was equilibrated for 1 ns, followed by a production run of 200 ns for each λ point. To prevent singularities caused by solute-solvent particle overlap as interactions were switched, a soft-core potential, as implemented in GROMACS, was applied (60). The soft-core parameters used were α=0.5 and p=1 (soft-core power). The free energies and their associated errors were calculated using the Multistate Bennett Acceptance Ratio (MBAR) method (61). The transfer free energy for moving a solute between two solvents, S1 and S2, *ΔG*_*s*1→*s*2_, was determined as the difference:

*ΔG*_*s*1→*s*2_ = *ΔG*_*s*1→*vac*_, − *ΔG*_*s*1→*vac*_.The calculated free energies for solvent pairs water/2-butanol, water/octanol, and water/chloroform were compared to available experimental data.

### Base pairing PMF calculations

PMF calculations were performed to quantify the free energy of hydrogen bonded base pairing in the CG model. Umbrella sampling (62) was used, following the protocol established for the Martini 2 RNA force field. Each nucleotide pair was placed in a 5 × 5 × 5 nm^3^ box, with bases constrained to remain coplanar. A series of 0.05 nm-spaced umbrella windows (0.3–2.0 nm) was used along the reaction coordinate, with a harmonic restraint of 1000 kJ□mol^−1^□nm^−2^. Each window was simulated for 200 ns for CG and 20 ns for AA model. The bases were aligned so that the X-axis passed through the N1 atom of the purines and the N3 atom of the pyrimidines, and the base plane was set as the XY plane. Position restraints with a force constant of 500 kJ□mol^−1^□nm^−2^ were used in the Y and Z directions. PMF profiles were computed using the weighted histogram analysis method (WHAM) (63) as implemented in GROMACS’ *gmx wham*, and errors were estimated via bootstrapping.

### ENM analysis and correlation maps

ENM has been widely applied to proteins using a single atom per residue (64) and to RNA structures using multiple atoms per nucleotide (65). For the analysis of the ADAT2/3-tRNA complex, we employed a model that represents the protein using alpha carbons (Cα) and the tRNA using its phosphate and ribose sugar backbone atoms (P and C1’), with a uniform spring constant and a cutoff of 1.2 nm. Correlation map values are normalized RMS average of 3-by-3 *x, y* and *z* correlations per bead:

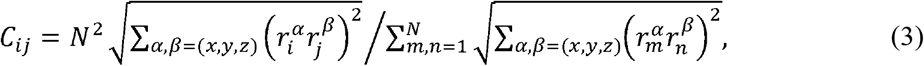

where 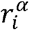 is a cartesian displacement component of bead *i* relative to its equilibrium position.

## Author Contributions

D.Y. and S.B.O. designed the research and prepared the manuscript. D.Y. performed the parameter development, simulations, validation and analysis. Funding acquisition: S.B.O.

## Acknowledgements

This work was funded by Bill & Melinda Gates Foundation INV-067331 and the National Institute of Health (grant no. GM147635 to S.B.O.) We also thank A. C. Vaiana for sharing all-atom MD trajectories of the ribosome complex.

## Data Availability

The code and tutorials for the MARTINI 3 Model for RNA are available on GitHub at https://github.com/DanYev/reForge as part of the package. A specialized version will be available for public access following peer review or upon request from the authors.

