## Supplemental Material for "Coarse-Grained RNA Model for the Martini 3 Force Field"

**Contents:** Supporting Figures S1–S3

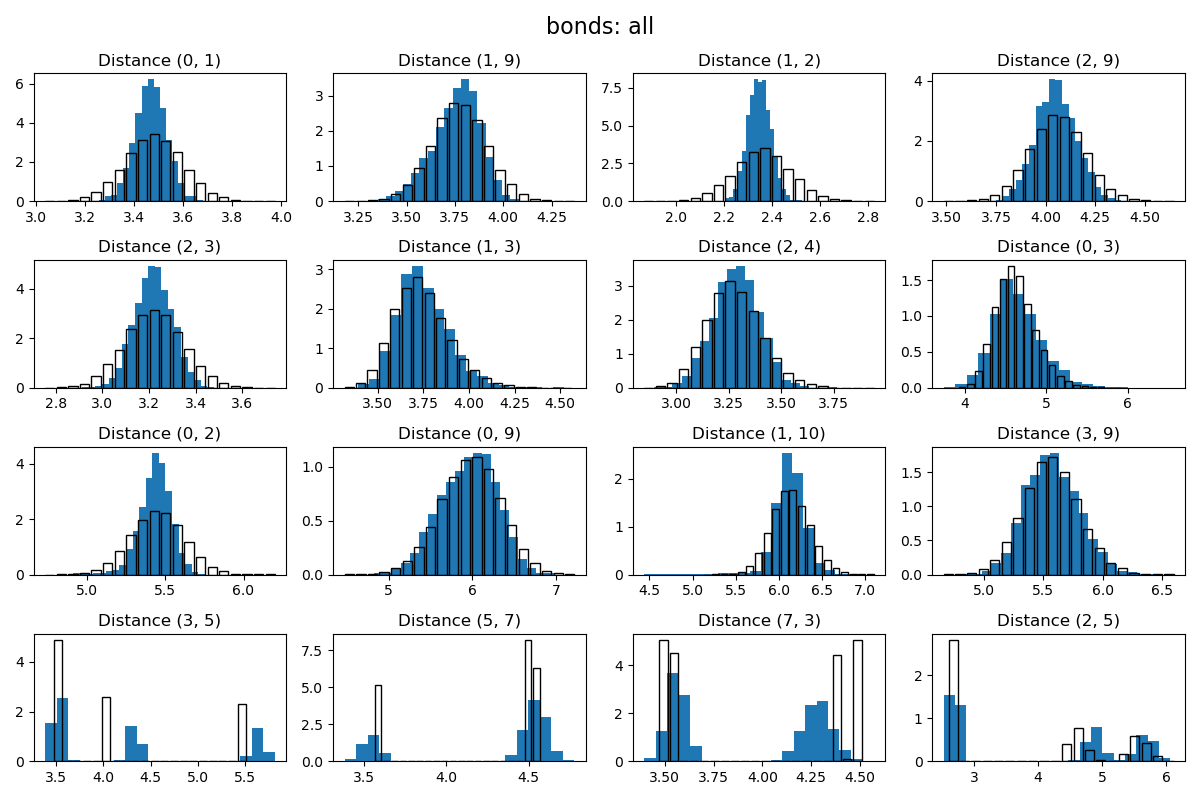

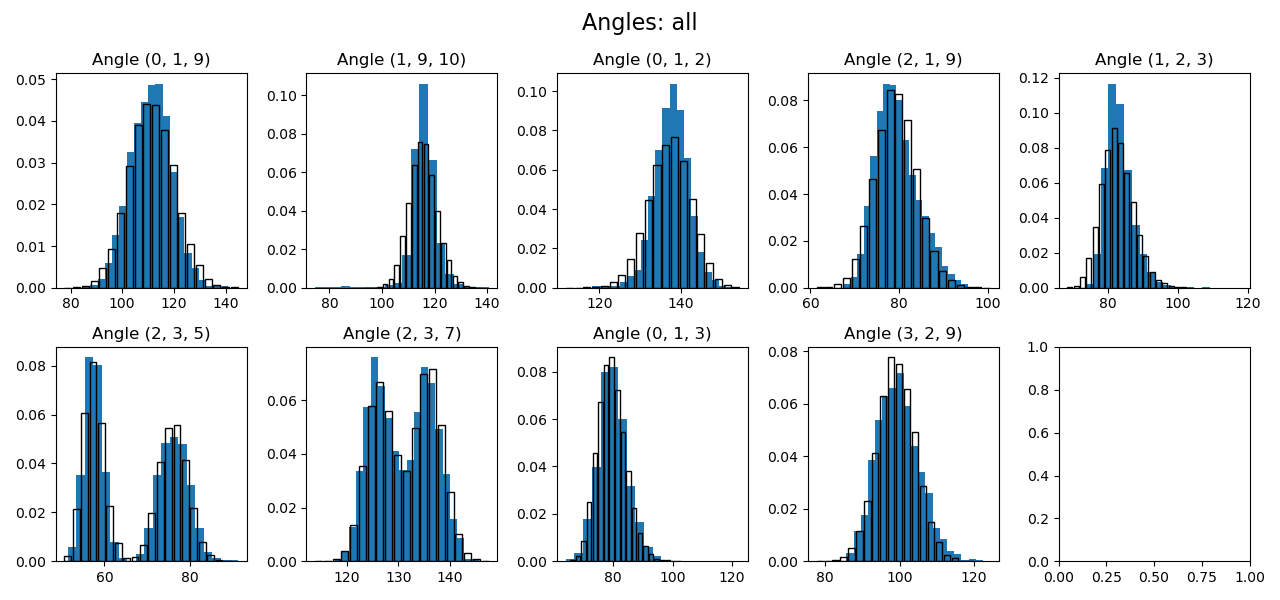

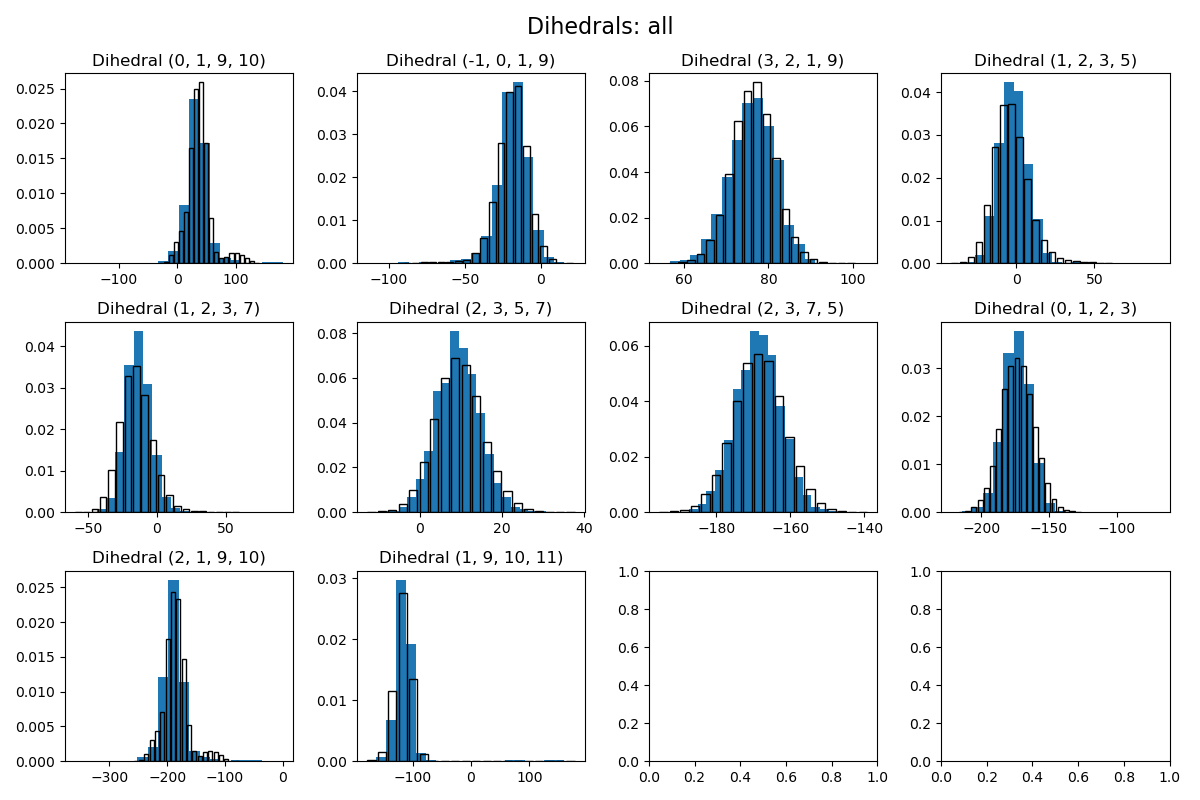

Figure S1. Distance, angle and dihedral distributions for double-stranded RNA

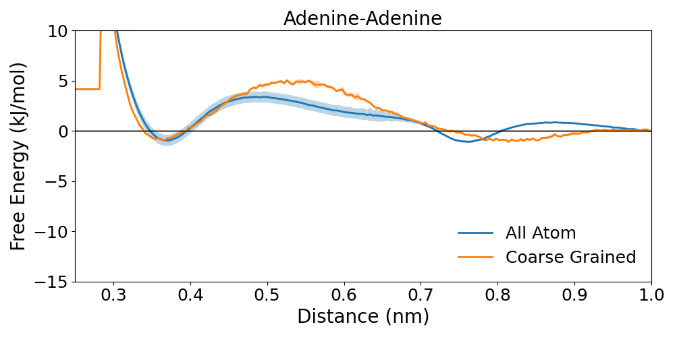

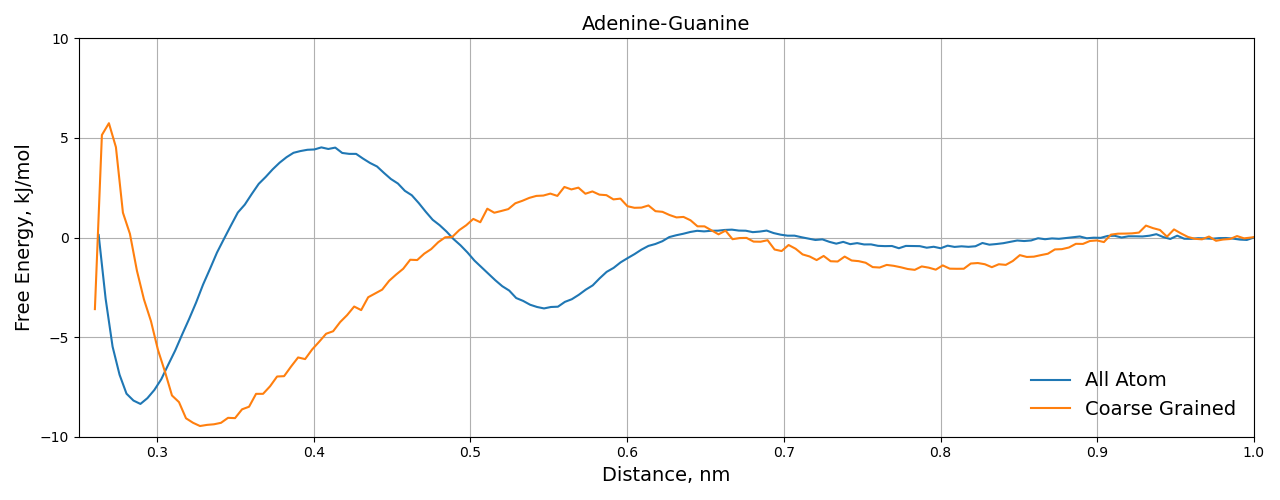

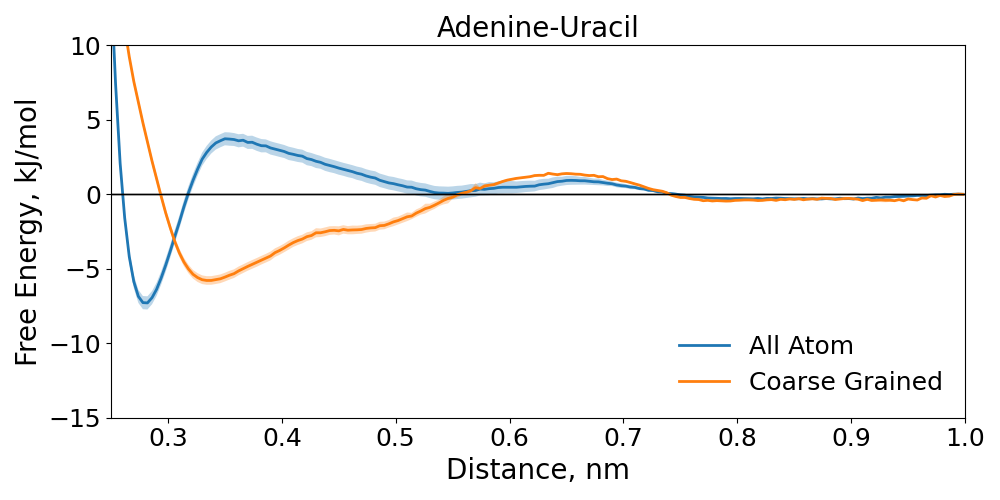

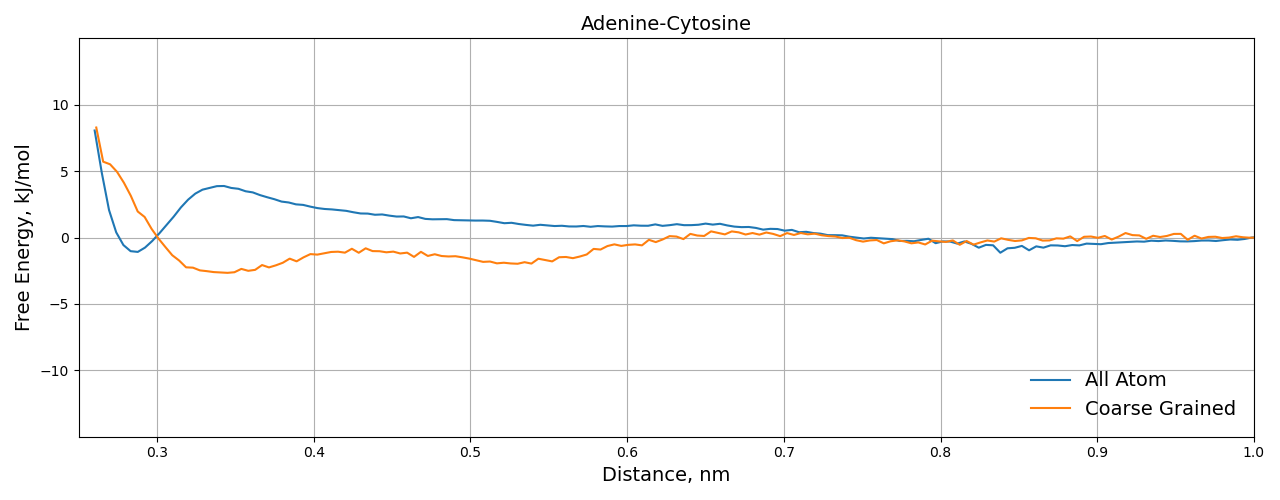

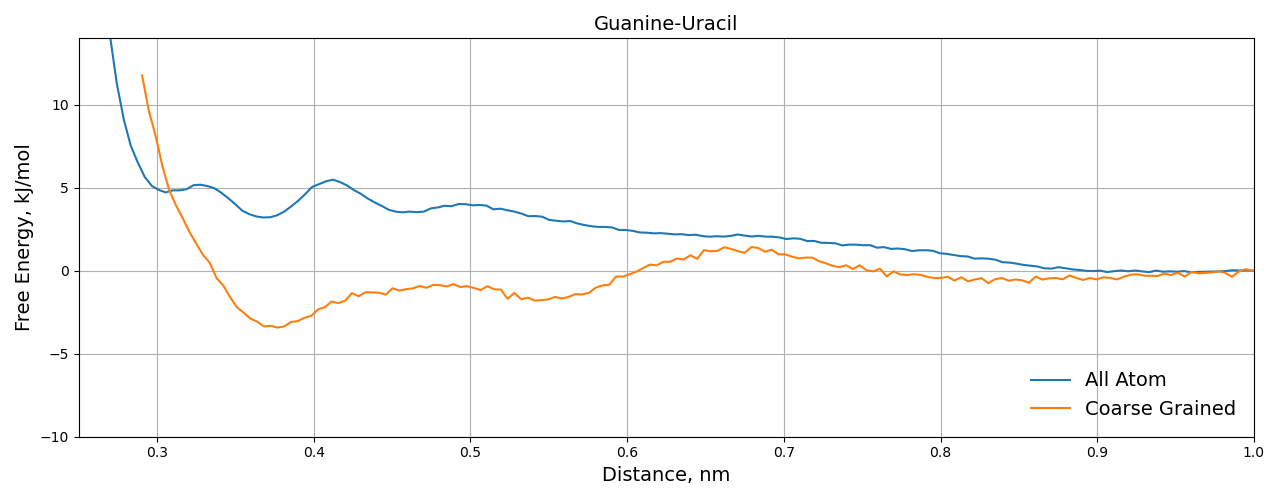

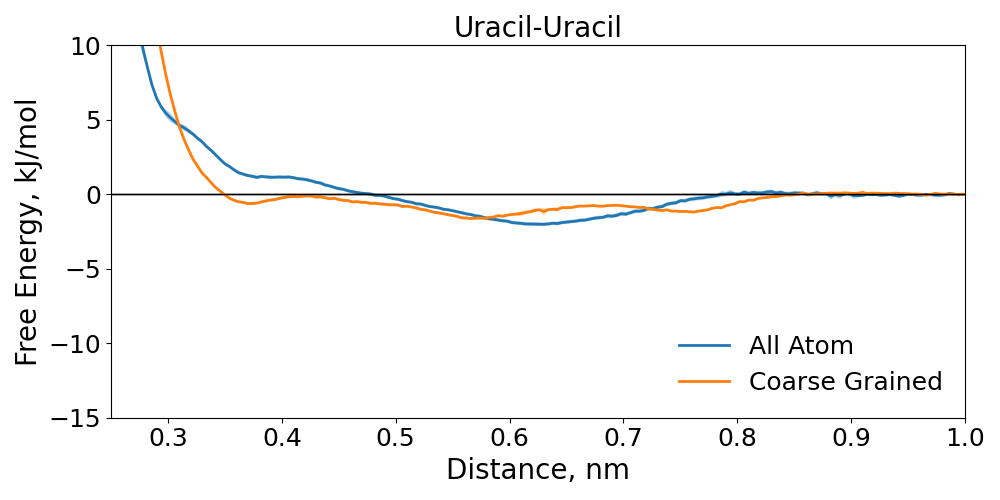

Figure S2. Dimerization PMF for the rest of the nucleobases

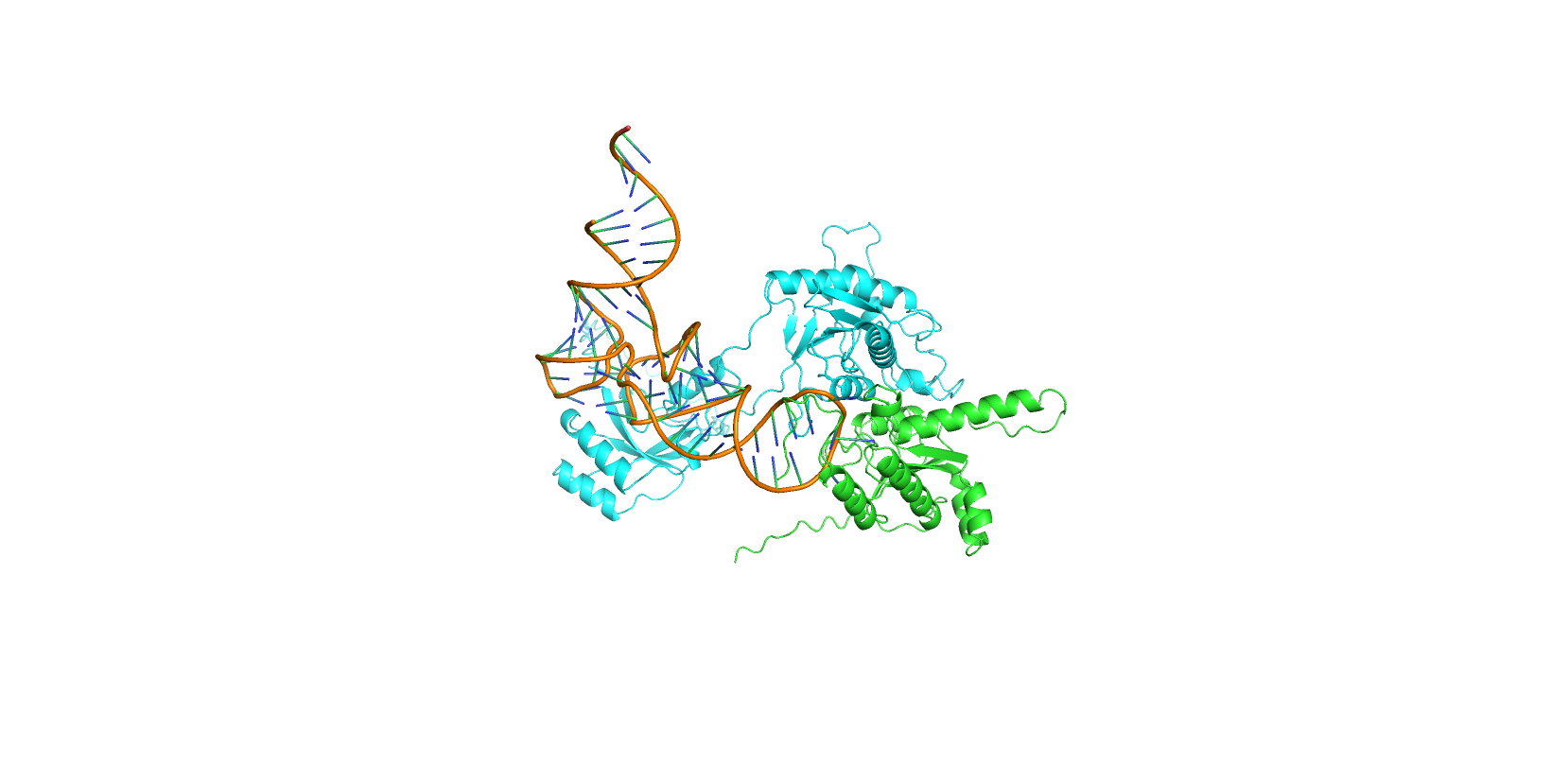

Figure S3. Structure of ADAT2/3 deaminase bound to a full-length tRNA.
